# Experimental *Trypanosoma cruzi* infection cured with a mechanistically distinct drug combination

**DOI:** 10.64898/2026.01.27.701990

**Authors:** Amanda Fortes Francisco, Francisco Olmo, Fanny Escudié, Eric Chatelain, John M. Kelly

## Abstract

Infections with the protozoan parasite *Trypanosoma cruzi* are widespread in the Americas and can lead to severe cardiac and/or gastrointestinal pathology. Current treatments are limited to monotherapies characterised by prolonged dosing regimens, disputed efficacy and toxic side-effects. Here, we demonstrate that short duration co-administration of well-tolerated sub-efficacious oral doses of the parasite-selective proteasome inhibitor GNF6702 and the pro-drug benznidazole produce parasitological cure in an experimental model of chronic Chagas disease.

---

Combination therapy has played a key role in controlling infectious diseases. With TB, malaria and HIV infection, it has had a major impact by reducing treatment failures and minimising resistance (1-3). However, this approach has yet to be applied to infections with the insect-transmitted hemoflagellate *Trypanosoma cruzi*. In the Americas, where >7 million people are affected, Chagas disease is the most serious parasitic infection (4,5). In addition, due to migration it has become a global health challenge (6). Infections are typically life-long and can lead to severe cardiac and gastrointestinal pathology, often with fatal outcomes (7,8). The only approved treatments are benznidazole (BZ) or nifurtimox, nitroheterocyclic pro-drugs that share the same bioactivation mechanism (9). In both cases, treatment is administered over prolonged periods (60-90 days), has variable efficacy, and is associated with adverse side-effects (10,11), highlighting an urgent need for innovative therapeutic strategies.

GNF6702 is a triazolopyrimidine-based selective inhibitor that targets the proteasome in kinetoplastid protozoa (12), and is well-tolerated in mice. The ubiquitin-proteasome system plays a pivotal role in protein homeostasis by degrading misfolded or damaged proteins and is essential for parasite survival and differentiation. GNF6702 exhibits a non-competitive mechanism of action, and binds allosterically to the parasite proteasome, inhibiting its chymotrypsin-like activity. Oral treatment of mice with GNF6702 was reported to reduce *T. cruzi* levels in the blood, heart, and colon to undetectable levels (12). However, as shown here (Fig. 1,2), GNF6702 monotherapy, either in vitro or in vivo, does not result in parasite elimination.

**Fig. 1.**
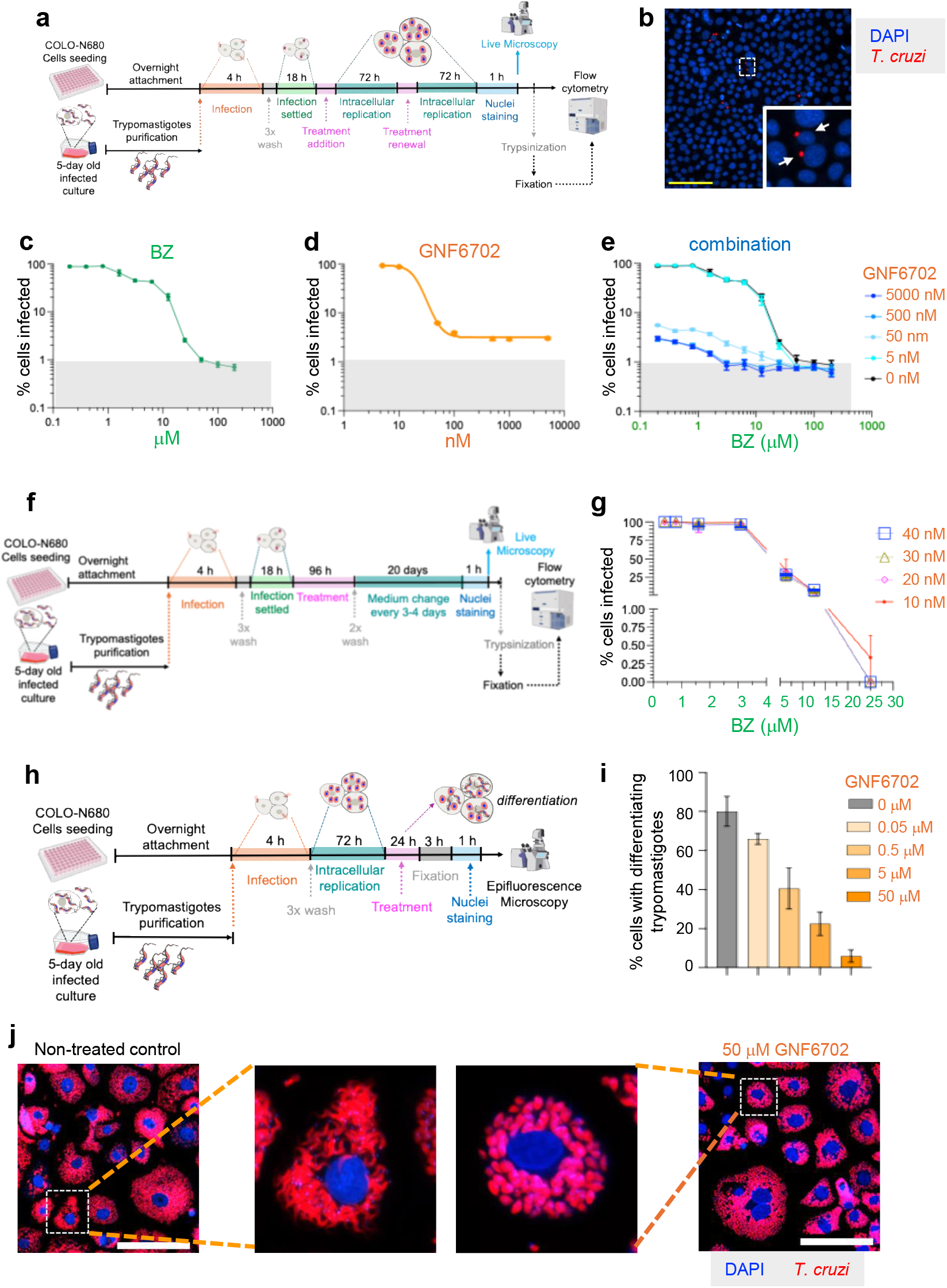
In vitro activity of GNF6702 and BZ. **a**, Protocol to determine intracellular activity. **b**, Fluorescence image of *T. cruzi* CL Brener PpyRE9h:mScarlet amastigotes (red) and host cell DNA (blue) following treatment with 6.3 μM BZ and 50 nM GNF6702, prior to flow cytometry. Yellow size bar=100 μM. Zoomed image of cells containing single amastigotes. **c-e**, Dose-response of BZ, GNF6702 and combination treatment after flow cytometry. Grey area indicates infectivity <1%. Data are the mean±SD (two independent biological replicates, three technical replicates). **f**, Protocol of “wash-out” assay to test combination treatment. **g**, Dose-response of combination treatment 20 days after drug removal measured by flow cytometry. **h**, Protocol to test impact of GNF6702 on amastigote to trypomastigote differentiation. **i**, Effect of GNF6702 on differentiation. Data (mean±SD) were derived from 3 independent wells in which 60-80 infected cells were inspected. **j**, Broad-field view of treated and non-treated cultures 96 hours post-infection. Zoomed images show single infected cells. Scale bars=100 µm.

**Fig. 2.**
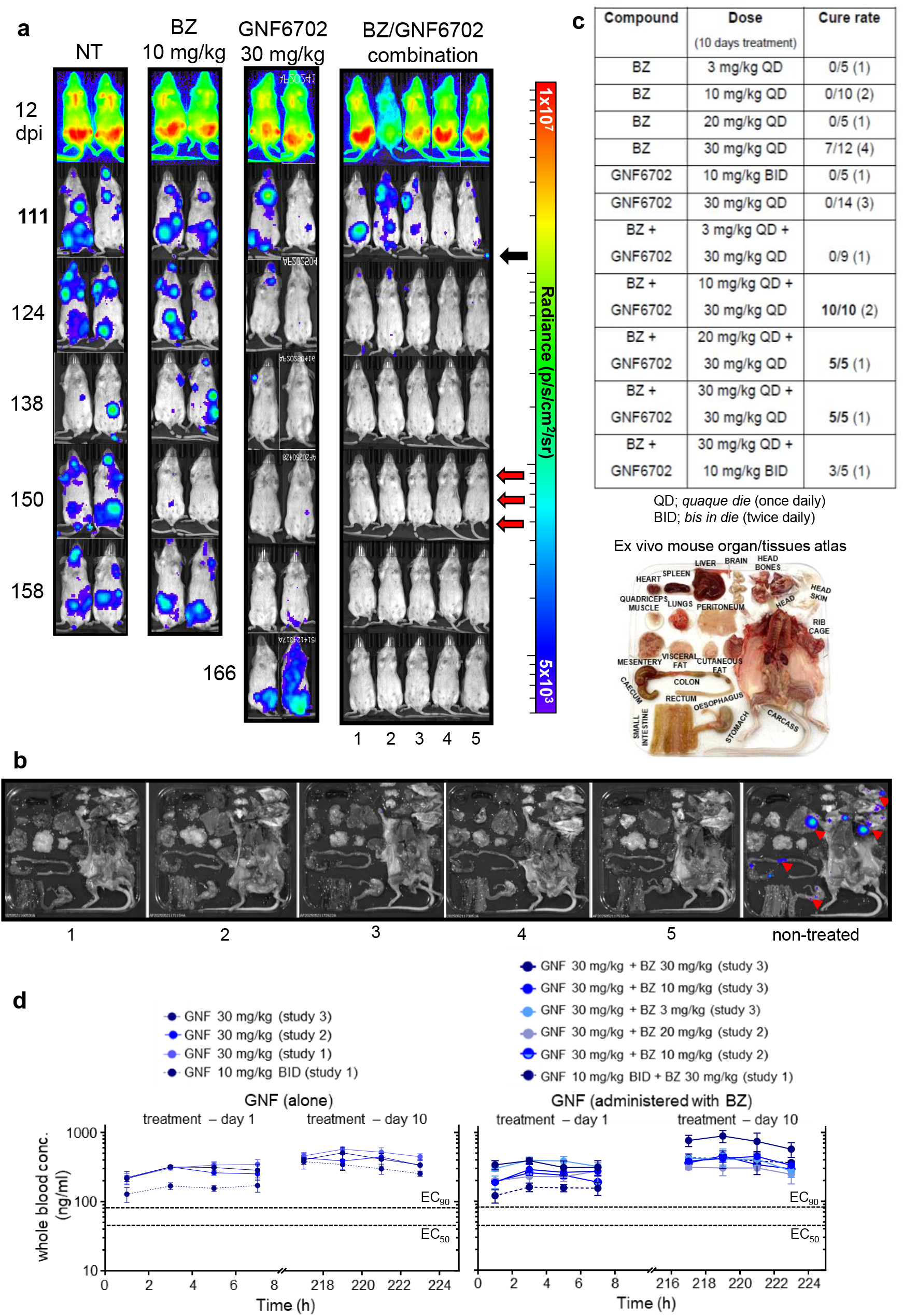
Efficacy of GNF6702:benznidazole combination therapy against chronic *T. cruzi* infection. **a**, Ventral images of BALB/c mice infected with CL Brener PpyRE9h:mScarlet strain (21), treated orally, once daily, for 10 days with 10 mg/kg benznidazole (BZ) and/or 30 mg/kg GNF6702. Treatment initiated 115 days post-infection (dpi) (black arrow). NT, non-treated. Mice were immunosuppressed using cyclophosphamide (3x200 mg/kg i.p.), administered at 3-day intervals (red arrows) starting 150 dpi. Heat-maps are on log10 scales and indicate bioluminescence intensity from low (blue) to high (red). **b**, Ex vivo imaging of GNF6702:BZ treated mice (Methods). Organs and tissues arranged as indicated in atlas. Bioluminescent foci are highlighted in non-treated image (right). **c**, Total cure-rates. Figures in brackets correspond to number of independent experiments from which data were derived. **d**, Pharmacokinetic profile of GNF6702 co-administered with BZ. Blood samples were taken from tail veins of infected mice on days 1 and 10 of treatment with GNF6702 alone or in combination with BZ. Whole blood concentrations were quantified using LC-MS/MS (Methods). Dotted lines identify EC_50_ (20.5 nM) and EC_90_ (36.9 nM) values.

In *T. cruzi*, BZ undergoes reductive metabolism which produces reactive intermediates (10,13), leading to thiol-depletion and damage to DNA, lipids and proteins. We hypothesised that combination therapy, based on a proteasome inhibitor (GNF6702) and a drug that can generate mis-folded/damaged proteins (BZ), might result in enhanced efficacy. To investigate this, we first assessed the effect on intracellular amastigote forms of the parasite (Fig. 1a,b). With BZ alone, 6-day treatment at concentrations >50 µM was required to reduce the number of cells infected with *T. cruzi* to <1% (Fig. 1c). Parasites that survive under these conditions are in a transient non-replicative state (14), and host cells that remain infected generally contain only a single parasite (as in Fig. 1b). An ability to eliminate such persisters will be a requirement for any new Chagas disease therapeutics. When tested alone, GNF6702 concentrations >25 nM led to a dramatic reduction in the number of infected cells, however complete parasite clearance could not be achieved, even at concentrations up to 5 μM. At all concentrations >100 nM, the infection persisted in at least 3% of cells (Fig. 1d). Combination treatment revealed highly potent activity. In the presence of 50 nM GNF6702, 3-4 times less BZ was required to reduce infected cells to <1% (Fig. 1b,e). Furthermore, exposure to 30 nM GNF6702 and 25 µM BZ was sufficient to eliminate intracellular parasites, when assessed 20 days after drug removal (Fig. 1f,g; Extended Data Fig. 1).

Trypomastigotes, the extracellular, non-replicative, infectious form of *T. cruzi* often display reduced drug susceptibility (15). In line with this, 4 hours pre-treatment of trypomastigotes with GNF6702 at doses up to 2 mM had no significant effect on infectivity (Extended Data Fig. 2a-c). To investigate the effect on differentiation of intracellular amastigotes to flagellated trypomastigotes, cultures were treated for 24 hours late in the infection cycle (72 hours) (Fig. 1h). This revealed concentration-dependent arrested transition of amastigotes into trypomastigotes (Fig. 1i,j), identifying a potential second means by which GNF6702 could act to enhance BZ potency. Amastigotes are the stage of the life-cycle most susceptible to BZ, whereas trypomastigotes are the least susceptible (15). GNF6702-mediated proteasome inhibition (12) therefore blocks differentiation into trypomastigotes, a life-cycle stage where higher BZ exposure would be required to eliminate the parasite.

In vivo efficacy of BZ/GNF6702 combination therapy was assessed using highly sensitive bioluminescence imaging (BLI) of infected BALB/c mice (16), a model widely used for Chagas disease drug assessment (17,18). As predicted from in vitro analysis (Fig. 1d), GNF6702 monotherapy (10-day oral dosing) failed to cure chronically infected mice (Fig. 2a), despite an initial knockdown in parasite burden. A similar outcome was observed during acute stage infections, even when treatment was prolonged to 20 days (Extended Data Fig. 3a). With BZ monotherapy, dosing at 30 mg/kg resulted in a partial cure-rate of chronic infections (10-day treatment), but lower doses were ineffective (Fig. 2c). To assess combination therapy, we first treated mice with GNF6702 at 10 mg/kg (10-days, twice daily), in combination with 30 mg/kg BZ (once daily). This produced a partial cure-rate (3/5). Next, we fixed GNF6702 at 30 mg/kg (once daily) and varied the BZ dose. At 10 mg/kg BZ and above, no parasites were detected in any mouse (n=20) following 10-day treatment (Fig. 2a-c). Curative outcomes were confirmed by post-treatment immunosuppression and ex vivo imaging of tissues and organs (19) (Fig. 2b). All treatment regimens tested were well-tolerated.

The failure of GNF6702 monotherapy to eliminate parasites under the conditions tested (Fig. 1c) was not due to insufficient exposure levels. Plasma concentrations exceeded the free EC_90_ of this compound (fraction unbound in plasma=0.063) across all treatment regimens tested, with slight accumulation at day 10 compared with day 1 (Fig. 2d). Thus, sustained exposure above the EC_90_ (30 mg/kg QD or 10 mg/kg BID) is not sufficient to achieve parasite elimination. In contrast, when combined with sub-efficacious BZ doses, full parasite clearance can be achieved. Blood concentrations for both drugs (Fig. 2d; Extended Data Fig. 4) were consistent with previously reported values (12,20). Importantly, GNF6702/BZ co-administration did not alter exposure of either compound. Collectively, these findings suggest that parasitological cure mediated by combination therapy results from the complementary modes of action of BZ and GNF6702 in preventing parasite persistence, rather than from pharmacokinetic interactions.

In summary, 10-day oral co-administration of the proteasome inhibitor GNF6702 (30 mg/kg) with low doses of BZ (10 mg/kg) results in a 100% cure-rate in a well-validated model of Chagas disease, with no adverse side-effects. By implication, combination therapy with these drugs, for which distinct modes of action have been established (9,12), eliminates the small number of persister parasites that frequently confound sterile clearance. Such short-duration therapy with low doses of BZ, in combination with GNF6702, should reduce adverse side-effects and improve compliance compared with the current therapeutic regimens. This represents a promising advance for treatment of a major Neglected Disease for which there have been no newly approved therapeutics for >50 years.

## METHODS

### Parasite/cell culture

Bioluminescent-fluorescent *T. cruzi* parasites (CL Brener PpyRE9h:mScarlet strain) and COLO-N680 cells (human oesophageal squamous cell carcinoma) were cultured in supplemented RPMI-1640 and Minimum Essential Medium (MEM, Sigma) respectively, as described (16,21). For infections, tissue culture trypomastigotes (TCTs) were derived from infected cells and exposed to cell monolayers for 4 hours. Extracellular parasites were then removed by washing with PBS, and flasks incubated with fresh medium for a further 5-7 days. Extracellular trypomastigotes were isolated by centrifugation of culture medium (1,600 *g*). Pellets were re-suspended in high-glucose (4.5 g/L) DMEM with 5% FBS and maintained at 37ºC until use. Motile trypomastigotes were counted using a haemocytometer.

### In vitro activity against *T. cruzi*

BZ and GNF6702 in vitro potency was determined by generating multiple-point potency curves by serial dilution in the corresponding culture medium. For amastigote assays, COLO-N680 cells in 100 µL growth medium were added to black, clear-bottomed, 96-well polystyrene microplates at 2.5×10^4^ cells/well. After overnight incubation, cells were infected with 5×10^5^ TCTs/well, a multiplicity of infection (MOI) of 10. Wells were then washed with PBS to remove non-internalized trypomastigotes, before adding 100 µL MEM supplemented with 5% FBS. Infections were allowed to establish overnight, then 100 µL MEM containing different drug concentrations was added. For assessment of activity against amastigote replication, 72-96 hours post-incubation, plates were washed with PBS and stained with 2 µg/mL of Hoechst for 1 hour. Cells were imaged by real-time epifluorescence microscopy using a Nikon Eclipse T2i. Cells were then washed with PBS and detached using TrypLE™ Express Enzyme at 37ºC, and fixed with 4% paraformaldehyde for 30 minutes. Cells were then washed in PBS, resuspended in flow cytometry staining buffer (PBS + 1% albumin) and fractionated in an Attune NxT Flow Cytometer. Gating was performed using a non-infected culture as a control. For trypomastigote assays, the protocol followed the same principle with modifications as indicated (Fig. 1h). Dose-response curves were fitted, and 95% confidence intervals calculated using the sigmoidal dose-response variable slope function from the Graph Pad Prism 10 software (www.graphpad.com). All experiments were performed twice, independently, with three technical replicates per concentration, unless otherwise stated.

### Mice and parasites

Animal infections were performed under UK Home Office project license PPL P9AEE04E4/PP7589959 and approved by the LSHTM Animal Welfare and Ethical Review Board. All protocols and procedures were conducted in accordance with the UK Animals (Scientific Procedures) Act 1986. Female BALB/c and CB17 SCID mice were purchased from Charles River (UK). Animals were maintained under specific pathogen-free conditions in individually ventilated cages. They experienced a 12-hour light/dark cycle, with access to food and water ad libitum. SCID mice were infected with 1×10^4^ TCTs in 0.2 mL D-PBS via i.p. injection. Female BALB/c mice, aged 7-8 weeks, were infected by i.p injection with 1×10^3^ blood trypomastigotes derived from a SCID mouse (19).

### Compounds and treatment

GNF6702 was synthesised by TCG, India and BZ by Epichem Pty Ltd., Australia. For in vitro assays, compounds were dissolved in DMSO (8 mM and 50 mM stock solutions), aliquoted and stored at -20ºC until required. For the in vivo studies, BZ and GNF6702 were formulated at different concentrations in 0.5% (w/v) hydroxypropyl methylcellulose (HPMC) and 0.4% (v/v) Tween 80 in Milli-Q H_2_O and 0.5% methylcellulose, respectively. Drugs and vehicle were administered by oral gavage according to weights at the beginning of treatment. For all studies, vehicle treated mice were used as the negative control, and BZ as the positive control.

### In vivo and ex vivo bioluminescence imaging

For in vivo BLI, mice were injected with 150 mg/kg d-luciferin i.p., anaesthetized using 2.5% (v/v) isoflurane in oxygen for 2-3 minutes, and imaged using an IVIS Spectrum system (Revvity, MA, USA) (16,19). Exposure times varied from 10 seconds to 5 minutes, depending on signal intensity. The detection threshold was established from uninfected mice. After imaging, mice were revived and returned to cages. For ex vivo imaging at the experimental end-points, mice were injected with 150 mg/kg d-luciferin i.p. 5 minutes before exsanguination under terminal anaesthesia using dolethal (200 mg/kg). Trans-cardiac perfusion was performed with 10 mL 0.3 mg/mL d-luciferin in DPBS (Dulbecco′s Phosphate Buffered Saline). Tissues and organs of interest were collected and arranged in squared Petri dishes, with the heart bisected along the coronal plane. Samples were soaked in DPBS containing 0.3 mg/mL d-luciferin. Bioluminescence imaging was performed as above. To estimate parasite burden, regions of interest were drawn using Living Image 4.8.2 to quantify bioluminescence expressed as total flux (photons/second) (16).

### Assessment of BZ and GNF6702 exposure in blood

Following dosing on day 1 and 10, blood samples (10 μL) were taken from the tail vein of infected mice that had been treated with BZ or GNF6702 at 1, 3, 5 and 7 hours, and transferred directly onto QIAcard FTA DMPK-B formats (QIAGEN; ref: WB129242) until analysis. Following extraction from the QIAcards, whole blood concentrations of BZ or GNF6702 were quantified using a validated LC-MS/MS method with a lower limit of quantification of 1.22 ng/mL.

Calibration and QC samples met acceptance criteria per internal guidelines. Non-compartmental analysis was performed using Phoenix WinNonlin software, version 8.1. GNF6702 total to free exposure was corrected using a fraction unbound (Fu) in mouse plasma of 0.063 and 0.05 in assay medium, as described (12). Similarly, a Fu of 0.68 in mouse plasma and 0.9 in assay medium was used for BZ.

### Statistical analyses

The statistical analysis was conducted using GraphPad Prism version 10.5.0. Groups were compared with ordinary one-way Anova with Tukey’s multiple comparisons test. A value of P <0.5 was considered significant.

## Acknowledgments

This research was supported by the UK Medical Research Council grants MR/T015969/1 to J.M.K. and funding from the Drugs for Neglected Diseases initiative (DNDi). DNDi received financial support for this work from the Federal Ministry of Research, Technology and Space (BMFTR) through KfW, Germany; Médecins sans Frontières, International; Swiss Agency for Development and Cooperation (SDC), Switzerland; and UK International Development, UK. FO was supported by Plan Propio of the University of Granada Research Stimulation grant (PP2023.PRI.I.14) and an institutional grant from the Fundación Ramón Areces (Ciencias de la vida y la materia). We would like to thank the Biological Services Facility team at LSHTM, especially James Gates and Carmen Abela for training, technical support, and scientific advice.

The authors declare no conflicts of interest.

All authors have read and agreed to the published version of the manuscript.

