## Extended data figures and legends for "Experimental *Trypanosoma cruzi* infection cured with a mechanistically distinct drug combination"

**Extended Data Fig. 1 | In vitro assessment of GNF6702:BZ combination therapy against amastigotes in COLO-N680 cells.** **a**, Broad field images illustrating the effect of GNF6702 treatment (6 days) on *T. cruzi* amastigote replication (CL Brener PpyRE9h:mScarlet strain). Scale bars=100 µm. **b**, Schematic of the combinational GNF6702:BZ "wash-out" assay. Each well was inspected exhaustively by epifluorescence microscopy to determine the presence of amastigotes, 20 days after drug removal. **c**, Broad field images of the wells in **b** at critical combination concentrations showing transition from "cure" to "non-cure" outcomes. Scale bars=50 µm.

**Extended Data Fig. 2 | In vitro inhibition of trypomastigote infection of COLO-N680 cells by GNF6702 and BZ.** **a**, Schematic protocol. **b**, Dose-response inhibition of COLO-N680 cell infection by *T. cruzi* trypomastigotes (CL Brener PpyRE9h:mScarlet strain) after 4 hours treatment with GNF6702 and BZ. The percentage infection was measured by flow cytometry. **c,** Broad field images of infected COLO-N680 cells 96 hours after invasion by drug-treated trypomastigotes (as above). Scale bars=50 µm.

**Extended Data Fig. 3 | In vivo efficacy of GNF6702 and BZ in acute and chronic murine models of *T. cruzi* infection.** BALB/c mice were infected with *T. cruzi* CL Brener PpyRE9h:mScarlet strain (21). Graphs show the total bioluminescence flux during infection and treatment of BALB/c mice (Methods) **a**, Acute stage monotherapy. **b**, Chronic stage monotherapy with BZ. The data relating to 30 mg/kg BZ were derived from one of four independent experiments. **c,** Chronic stage monotherapy with GNF6702. **d**, Chronic stage combination therapy. Grey and pink shading identifies the treatment and immunosuppression periods, respectively. The horizontal black line represents the background bioluminescence level, determined from non-infected mice.

**Extended Data Fig. 4 | Pharmacokinetic profile of BZ co-administered with GNF6702.** Blood samples were taken from the tail vein of infected mice on days 1 and 10 of treatment with BZ alone or in combination with GNF6702. Whole blood concentrations of BZ were quantified using LC-MS/MS (Methods).

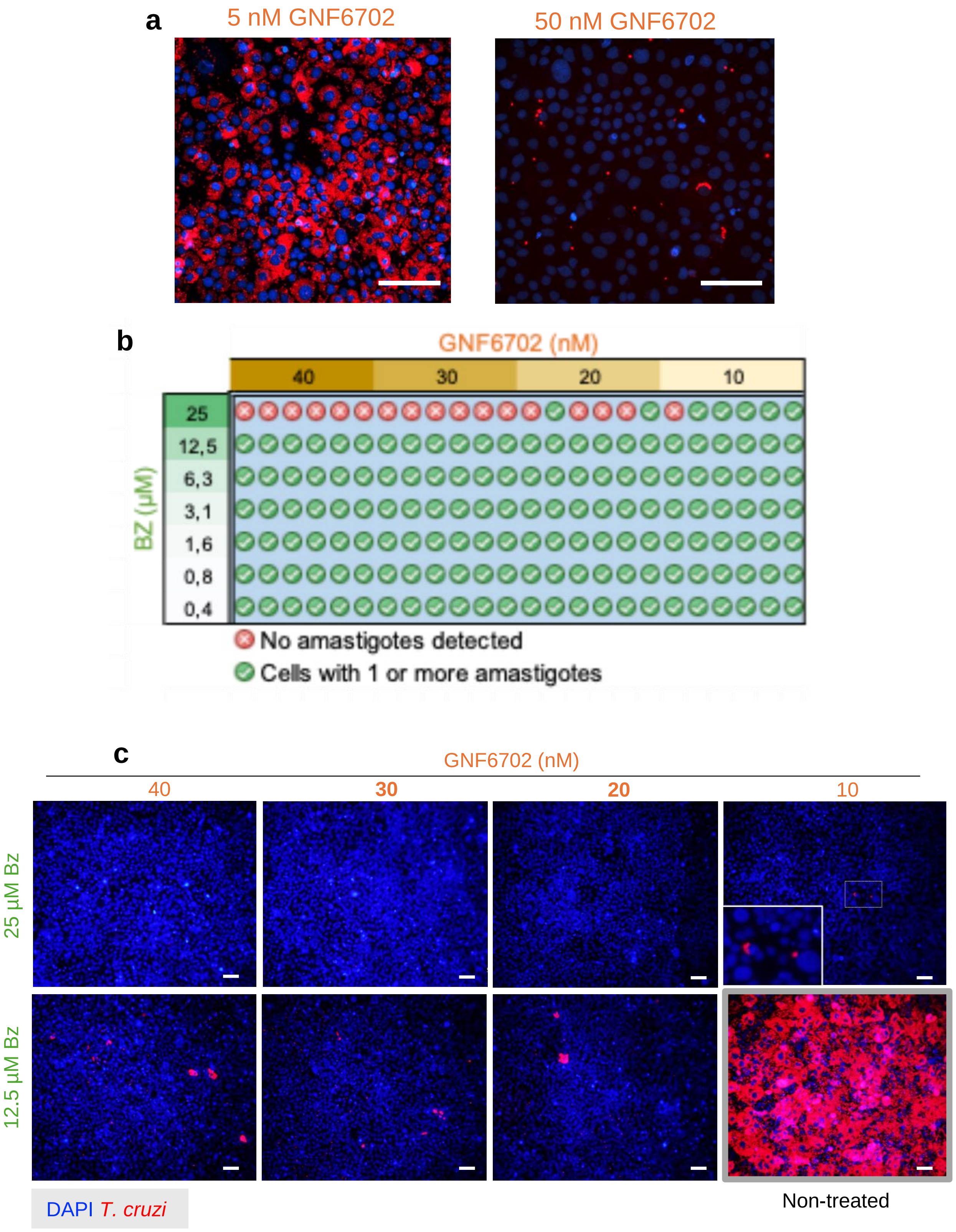


Extended Data Fig. 1

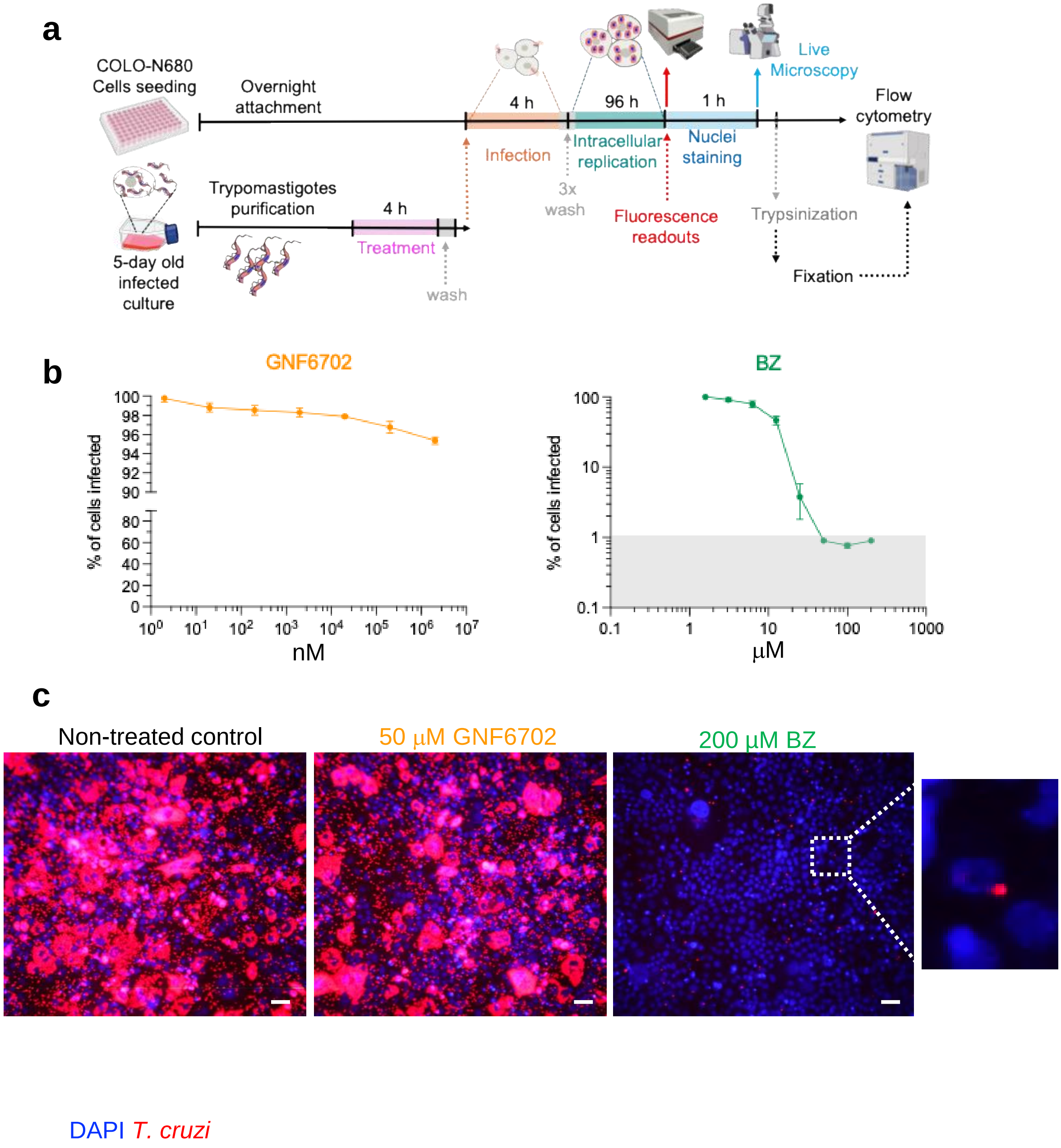


Extended Data Fig. 2

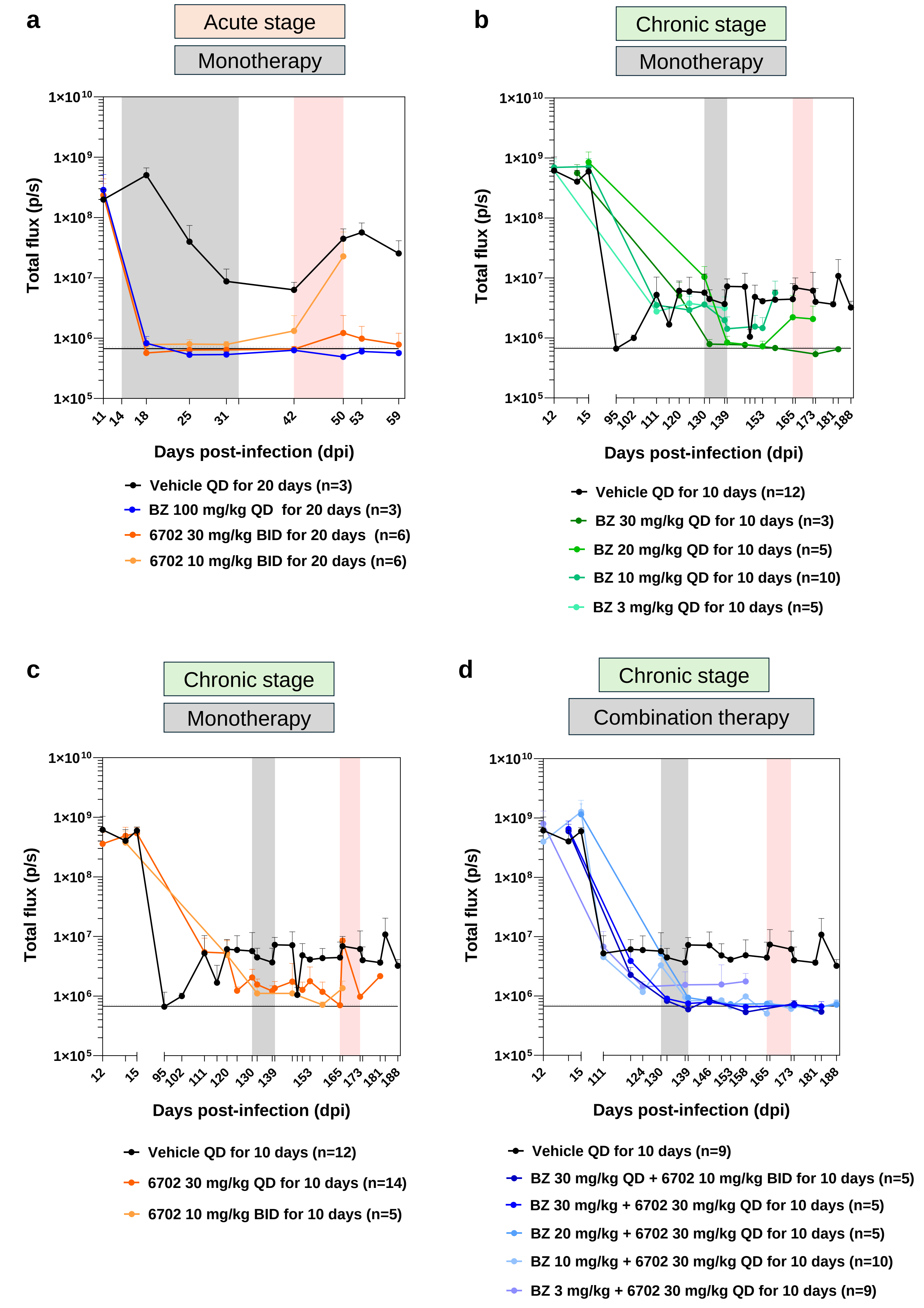


Extended Data Fig. 3

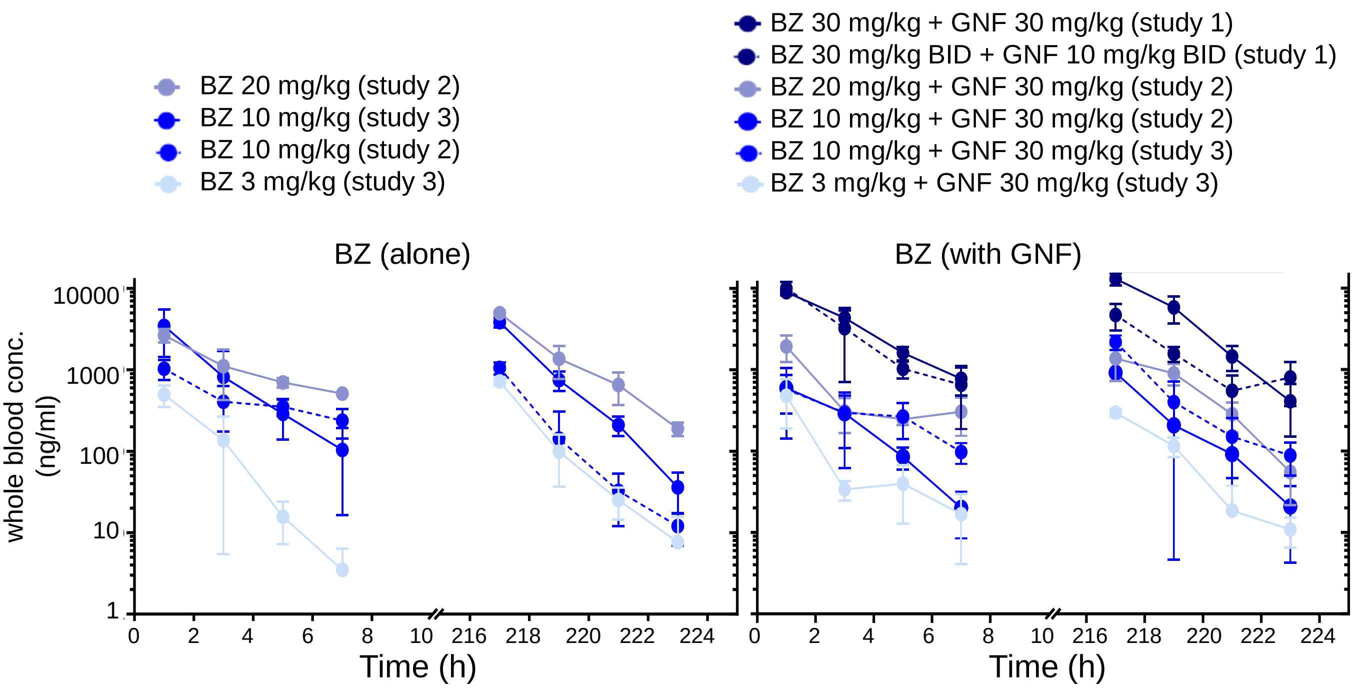


Extended Data Fig. 4
